# GRIBCG: A software for selection of sgRNAs in the design of balancer chromosomes

**DOI:** 10.1101/484360

**Authors:** Brian B. Merritt, Lily S. Cheung

## Abstract

**Background:** Balancer chromosomes are tools used by fruit fly geneticists to prevent meiotic recombination. Recently, CRISPR/Cas9 genome editing has been shown capable of generating inversions similar to the chromosomal rearrangements present in balancer chromosomes. Extending the benefits of balancer chromosomes to other multicellular organisms could significantly accelerate biomedical and plant genetics research.

**Results:** Here, we present GRIBCG (Guide RNA Identifier for Balancer Chromosome Generation), a tool for the rational design of balancer chromosomes. GRIBCG identifies single guide RNAs (sgRNAs) for use with *Streptococcus pyogenes* Cas9 (SpCas9). These sgRNAs would efficiently cut a chromosome multiple times while minimizing off-target cutting in the rest of the genome. We describe the performance of this tool on six model organisms and compare our results to two routinely used fruit fly balancer chromosomes.

**Conclusion:** GRIBCG is the first of its kind tool for the design of balancer chromosomes using CRISPR/Cas9. GRIBCG can accelerate genetics research by providing a fast, systematic and simple to use framework to induce chromosomal rearrangements.

## Background

Balancer chromosomes contain multiple inverted regions capable of suppressing crossovers during meiosis. They also contain dominant mutations that allow their unambiguous tracking during crosses, and recessive lethal mutations that prevent the recovery of homozygous progeny. These features make balancer chromosomes particularly useful in preventing the loss of recessive lethal or sterile mutations from a population (without manual selection) and during saturation mutagenesis screens [1–3]. In plant breeding, balancer chromosomes could help preserve the advantages of heterosis without full apomixis [4].

CRISPR/Cas9 genome editing can generate inverted regions similar to the rearrangements present in balancer chromosomes (Figure 1) [5, 6]. Chromosomal rearrangements have been reported in *C. elegans* and zebrafish germlines, and in pig, mouse, and human somatic cells [5, 7–10]. Most notably, CRISPR/Cas9 was used to generate a large inversion at a specific site in C. *Elegans;* in a part of the genome that was previously not covered by any balancer region [5].

**Figure 1.**
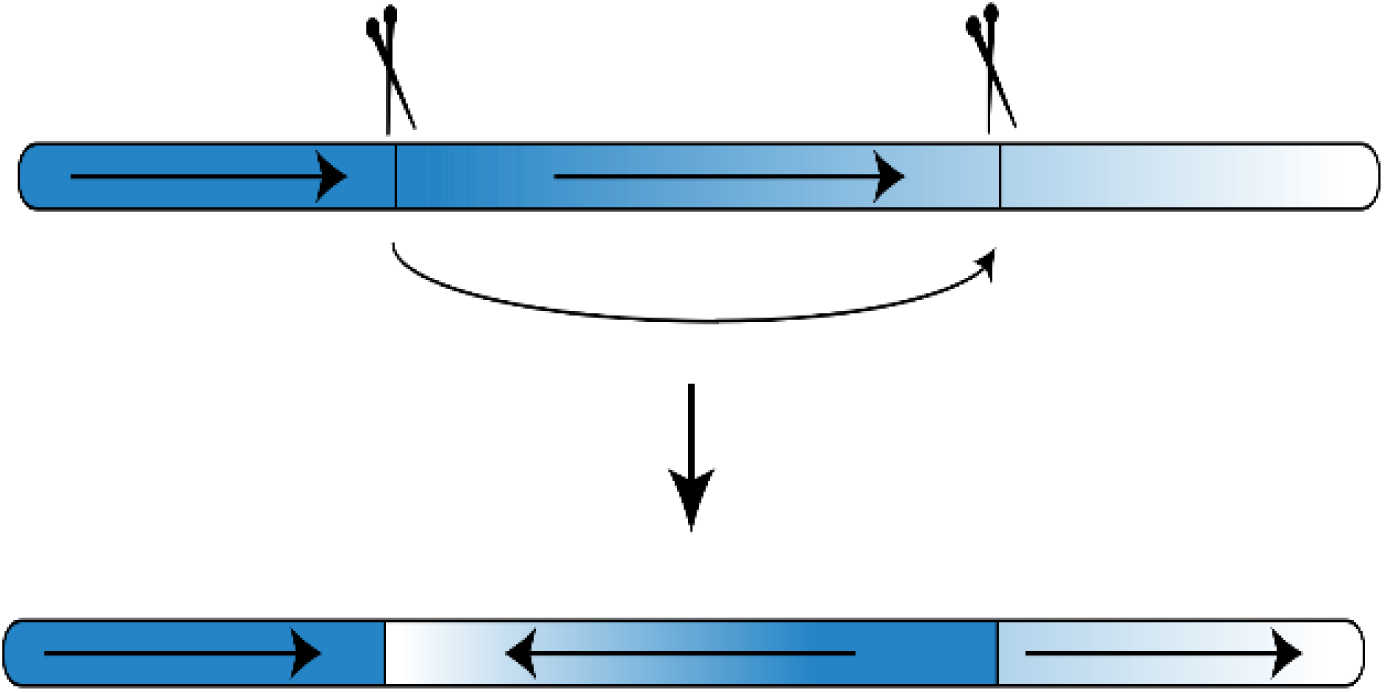
Double-stranded breaks in multiple sites along the same chromosome arm can result in inversions. CRISPR/Cas9 can be used to target specific regions within an arm.

The Cas9 complex consists of two primary components, the SpCas9 enzyme from *S. pyogenes* [6] and a single guide RNA (sgRNA). Each sgRNA consists of a 20-bp spacer sequence and an upstream 3-bp Protospacer Adjacent Motif (PAM) [6, 11]. Double-stranded breaks are induced by the annealing of the sgRNA to the target DNA then followed by Cas9 cutting [11]. Cells then repair this break via homology-directed repair (HDR) or non-homologous end-joining (NHEJ). Double-stranded breaks in multiple sites along the same chromosome can result in inversions [12].

The efficiency of Cas9 cutting is reduced by mismatches between the PAM or spacer sequence and the target DNA. Mismatches in the PAM are poorly tolerated [13]. As a result, sgRNAs with high potential of cutting by SpCas9 will primarily contain a 5’-NGG-3’ PAM, where N is any DNA nucleotide. Mismatches in the spacer sequence affect cutting efficiency in both a position and a nucleic identity dependent manner [13].

Multiple tools have been developed for the optimal design of sgRNAs. These tools account primarily for the thermodynamics of binding, secondary structure properties, and position-dependent nucleotide compositions [14, 15]. Thermodynamic considerations contributing to the on-target activity of sgRNAs include GC content, entropy change, enthalpy change, free energy change, and melting temperature [14]. Secondary structure features include repetitive sequence counts, length of potential stem-loops, minimum energy of folding, and the longest polyN for a sequence [14].

Here we describe GRIBCG (Guide RNA Identifier for Balancer Chromosome Generation), a tool to enable balancer chromosomes in multicellular organisms other than flies. GRBICG is a Perl and R based tool designed to be locally run on any computer. It is designed to accept any FASTA file containing a single genome and is freely available at https://sourceforge.net/p/gribcg/code/ci/master/tree/

GRIBCG identifies ideal sgRNAs for balancer chromosome generation based on on-target efficiency, off-target effects, and coverage. It selects sgRNAs that would cut a given chromosome multiple times, while minimizing off-target cuts in the rest of the genome. In *D. melanogaster*, it has been estimated that recombination events are suppressed within 2 Mbps on each side of an inversion breakpoint [1]. Our tool accounts for this fact by optimizing coverage, defined here as the percentage of a chromosome that is protected from recombination due to their proximity to an inversion breakpoint. Our choice of design parameters is intended to minimize the number of generations that must be screened in order to experimentally recover the balancer chromosomes.

Finally, to benchmark the precision and efficiency of our tool, we applied GRIBCG to several model organisms: mouseear cress (*A. thaliana*), fruit fly *(D. melanogaster)*, worm *(C. elegans*), zebrafish *(D. rerio)*, mouse *(M. musculus*), and rice (*O. Sativa)*, and successfully identified optimal sgRNAs with 70% or more coverage. We also compare the result of our tool with two routinely used *D. melanogaster* balancer chromosomes.

## Implementation

GRIBCG requires users to upload FASTA chromosome sequences. Additionally, GRIBCG can accept a FASTA file containing locations of all genes associated with a given organism. The pipeline selects ideal sgRNAs based on on-target, off-target, and coverage properties. GRIBCG is designed for local use in desktops or laptops, thus it is accessible through a graphical user interface (GUI). Users may upload FASTA-formatted files containing known gene start and stop locations for each chromosome. An overview of the pipeline is depicted in Figure 2 and the GUI in Figure 3.

**Figure 2.**
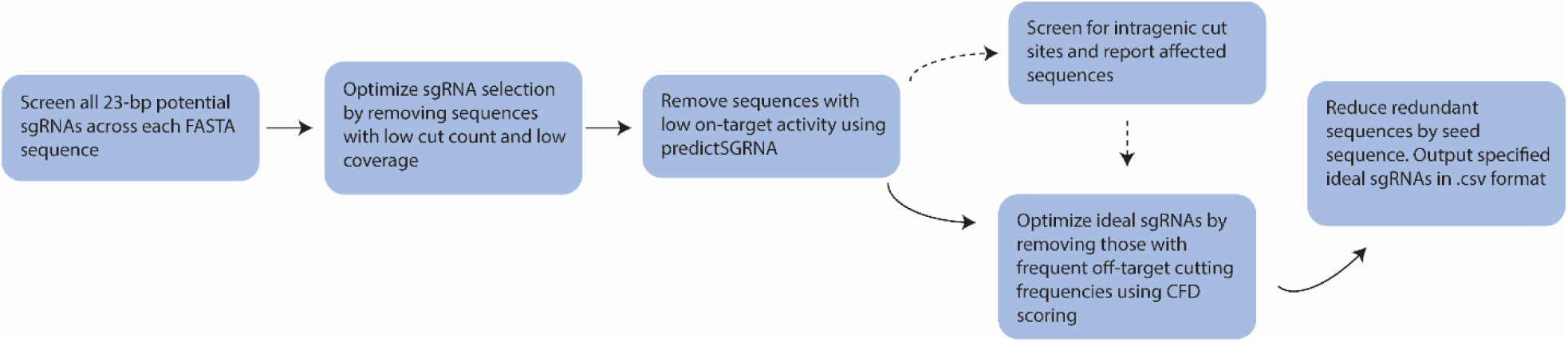
GRIBCG procedural steps in the generation of top sgRNA lists. Each chromosome is analyzed, selecting all potential 23-bp sequences containing 5’-NGG-3.’ Then, 18-bp partial sequence sites (PSS) are screened across all other chromosomes, producing an average cut distance for a given PTS based on PSSs. Off-target scoring calculated by counting presence of PSSs for given PTS on other chromosomes. PTSs with PSS sites on other chromosomes are rejected. Potential target sequences are then filtered by optimal cut counts for their chromosome. On-target scoring is performed for all remaining PTSs. A final list of top sgRNA designs per chromosome is generated based on off-target, on-target scoring, and coverage. Dashed arrows indicate optional parameters where users may upload a single file containing gene locations.

**Figure 3.**
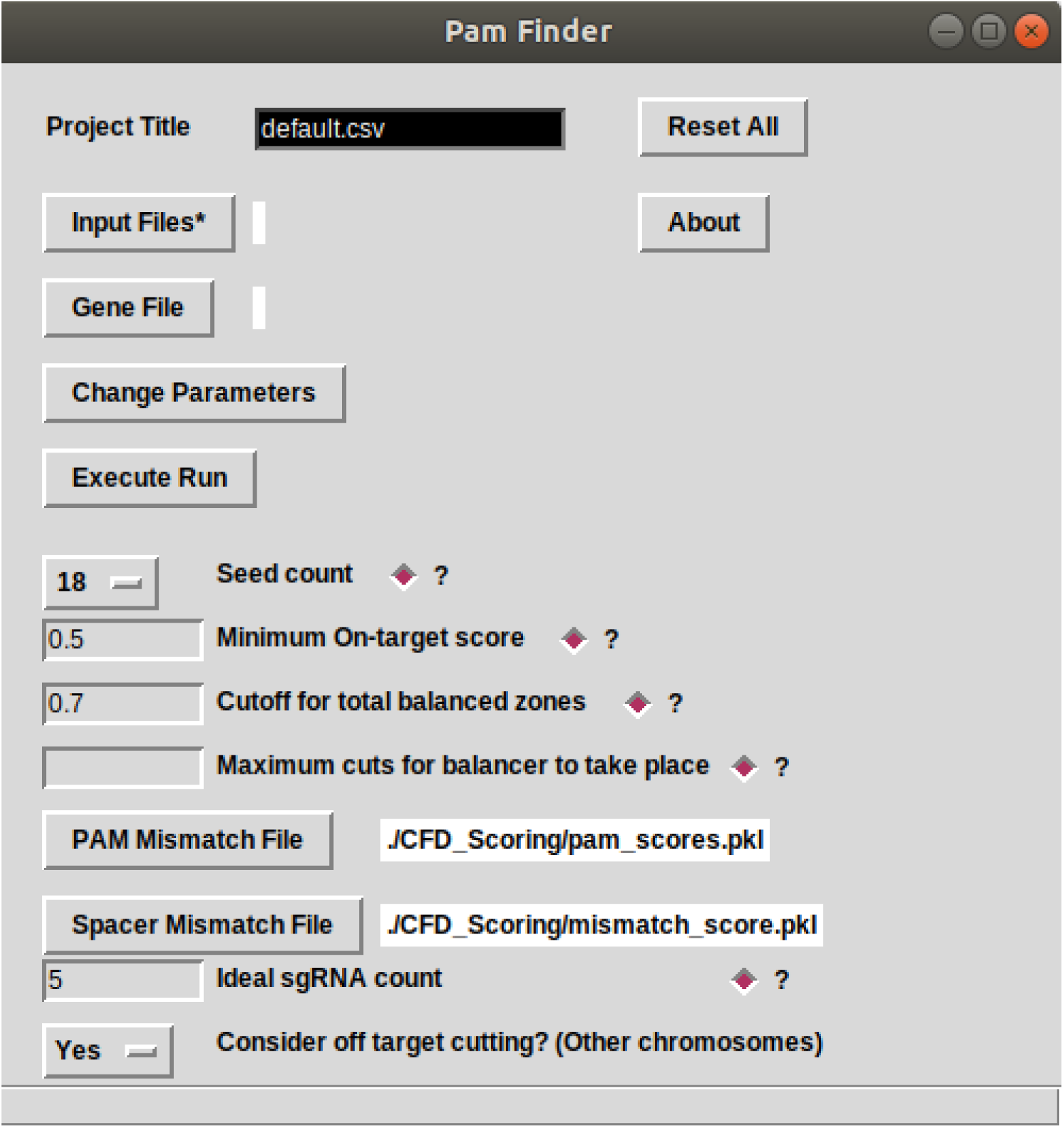
GUI depicting the different options available to the user in GRIBCG. The tool provides a way to change individual parameters when filtering potential guide RNAs. In addition, each parameter comes with a short description of its role in the design process.

First, GRIBCG searches for all potential sgRNA target sites in each chromosome. Bioperl is utilized for the sequence accession analysis. Each chromosome is analyzed for the presence of, on both strands, a given PAM (5’-NGG-3’ for *S. pyogenes Cas9*). The exact 23-bp potential target sequence (PTS) along with chromosomal position and flanking sequences is recorded. Each PTS has a corresponding partial sequence (PSS) or seed sequence. Due to the annealing properties of the CRISPR/Cas9 system to PTSs, nucleotide matched identity is weighted by their proximity to the PAM sequence (downstream). Due to the considerable size of many genomes, this tool often has variable performances in both computation cost and time. In order to limit memory usage, a temporary file is created containing all PTSs in the uploaded FASTA chromosome file(s). *M. Musculus*, for instance, required 19 GB of space during the generation of a single temporary file.

Next, potential sgRNAs are binned to reduce computation complexity. GRIBCG merges all cut locations for each binned group based on PTS sites. We perform this step to reduce computation complexity as comparing efficiency scores for sgRNAs yields a computation complexity of O(n^2^) and therefore can be up to 10^7^ unique sgRNAs in larger genomes. Our choice to use this seed sequence is validated by the experimentally determined effect of mismatches between positions from Hsu et al [16].

GRIBCG then analyzes total coverage of an entire chromosome based on Cas9-induced breakpoint positions. Considering a total of 4 Mbp surrounding a breakpoint, the algorithm calculates the ideal cut count and filters out PSSs bins that exceed this threshold. It is important to note that the distance between PTSs, and thus between potential breakpoints, often varies widely. For instance, PTSs may contain identical cut counts yet hold different coverages because of the proximity between sites (Figure 4).

**Figure 4.**
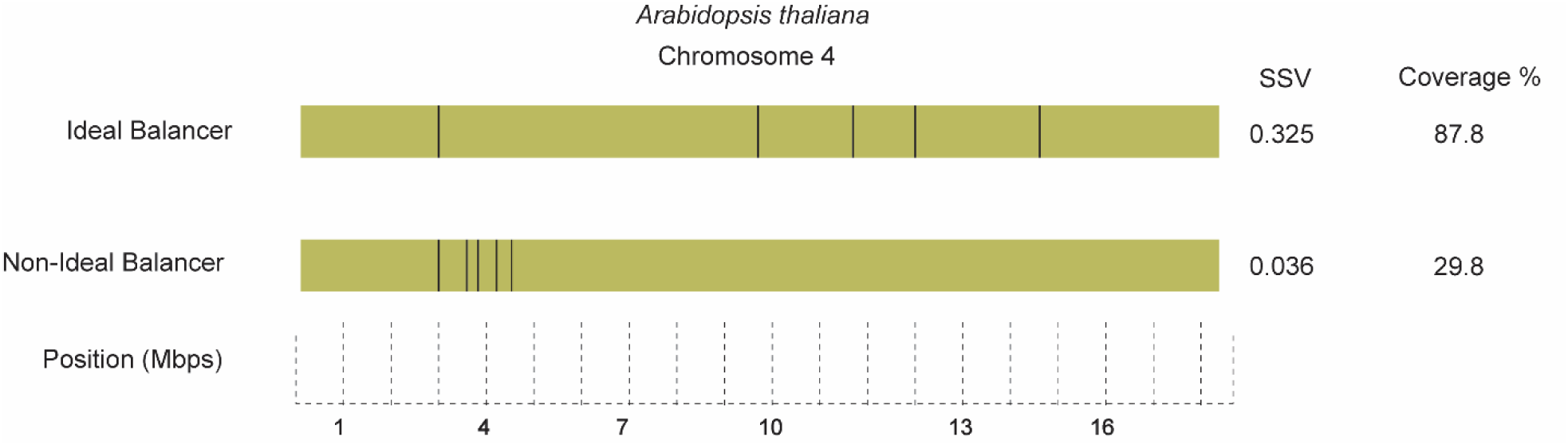
Ideal balancer (top) produced by a single sgRNA in fourth chromosome of *A. thaliana*. Each inversion breakpoint protects the surrounding 4 Mbps from recombination. GRIBCG optimizes coverage, resulting in more evenly spaced breakpoints. A non-ideal balancer (below) produced by a single sgRNA, on the same chromosome as above, where breakpoints are situated near one another leaving most of the chromosome unprotected. Each vertical line represents a given double-stranded break.

sgRNAs surpassing the predefined coverage threshold are then analyzed for on-target activity. All PTSs on-target scores are then calculated via the R-tool predictSGRNA [14, 15]. This tool analyzes PTS candidates based on property models from existing CRISPR datasets and provides a list of efficiency scores for each PTS. PSS bins with average on-target efficiencies less than the pre-defined on-target threshold are removed from further analysis.

GRIBCG calculates off-target activity to minimize undesired double-stranded breaks. For instance, mismatches between the sgRNA and the PTS has varying effects on Cas9 activity [16]. This leniency is accounted for by weighing base mismatches and assigning an off-target score. Due to the extensive filtering performed, the algorithm can afford to utilize a new mismatch analysis, the Cutting Frequency Determination (CFD) [12]. CFD considers both nucleic identity and position parameters as metric of determining the frequency of a cut based on mismatch percentage. Each mismatch is pooled into a product of penalty scores to give a CFD value between 0 (least efficient) and 1 (most efficient). This allows GRIBCG to determine undesirable off-target cuts for each PSS bin. Each PTS is then compared to all other PTSs on the remaining chromosomes in order to find probable off-target sites. For example, each PTS on the first chromosome would be compared to all PTSs not on the first chromosome. A total score is summed for each PTS and all probable off-target sites are reported.

Finally, GRIBCG defines a Simulated Sequence Value (SSV) as the final metric used to select the ideal sgRNAs. This metric is calculated by standardizing all PTSs on their respective chromosomes based on total chromosomal coverage and off-target efficiency scoring. By default, the tool considers both off-target and coverage features, but a user may opt to remove the consideration of off-target effects. The top sgRNAs (default of 5) are then reported with their corresponding SSV.

## Discussion and results

We implemented GRIBCG to generate sgRNAs for six of the best-established model organisms (Table 1). We present a case study of *A. thaliana*, which had 70% or more coverage for the top sgRNAs of each chromosome (Table 2). A total of 8,099,451 unique potential cut sites were screened. From there, all seed sequences were binned to give a total of 7,541,563 sequences. Thresholding of coverage further reduced the number of sequences to 2,804 multi-site cutting seed sequences. On-target efficiency was then analyzed on every target site and averaged across each sequence respective to their chromosome. After filtering, a total of 7,145 sites remained. Finally, the off-target frequency was analyzed using CFD scoring to optimize on-target and off-target cutting.

**Table 1.** Performance metrics of GRIBCG on various genomes

| Organism | User Time (s) | System Time (s) | Memory Peak (Mb) |
| --- | --- | --- | --- |
| *C. elegans* | 86.14 | 34.14 | 1042.5 |
| *A. thaliana* | 105.99 | 40.63 | 970.8 |
| *D. rerio* | 1419.52 | 478.39 | 2121.7 |
| *M. Muscula* | 3293.55 | 1601.28 | 8307.8 |
| *O. sativa* | 605.87 | 206.23 | 1969.2 |
| *D. melanogaster* | 175.68 | 62.66 | 2609.6 |

**Table 2.**
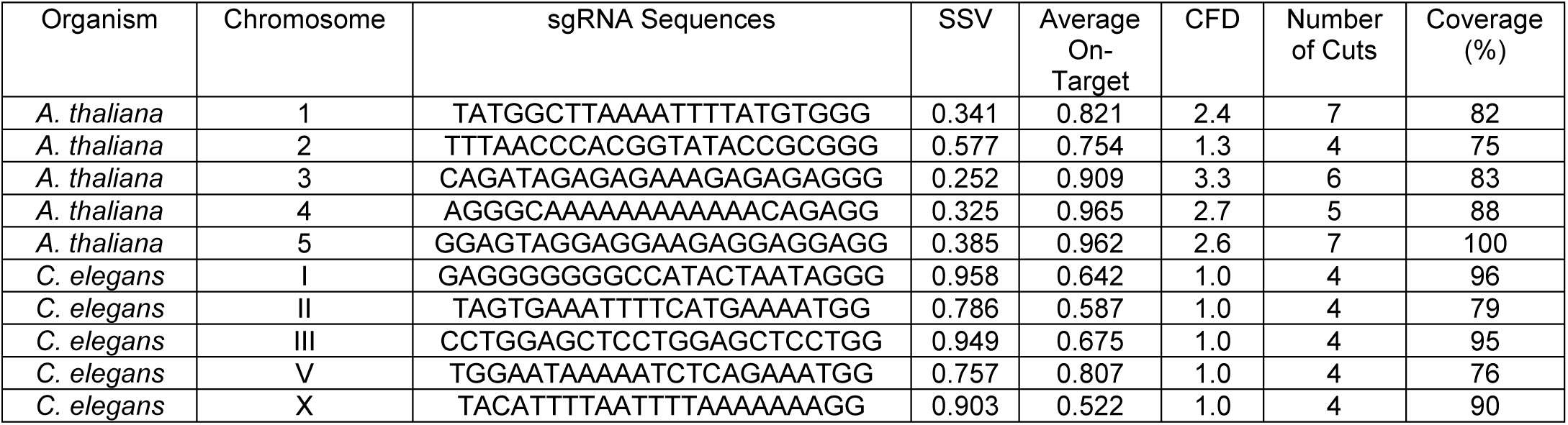
Top ideal GRIBCG-generated sgRNAs for each chromosome in *A. thaliana* and *C. elegans.*

We compared the results from GRIBCG to two of the most commonly used balancer chromosomes in *D. melanogaster.* Figure 5 depicts the locations of all potential inversion breakpoints throughout the second (SM6a) and third (TM3) balancer chromosomes in *D. melanogaster* [1]. The estimated coverage of these balancer chromosomes are 46% and 52%, respectively. In comparison, the top GRIBCG-selected sgRNAs that would result in the same number of breakpoints for the second and third chromosomes cover 57% and 61%, respectively. This suggests that newly generated balancer chromosomes designed with our tool would perform similarly to existing ones.

**Figure 5.**
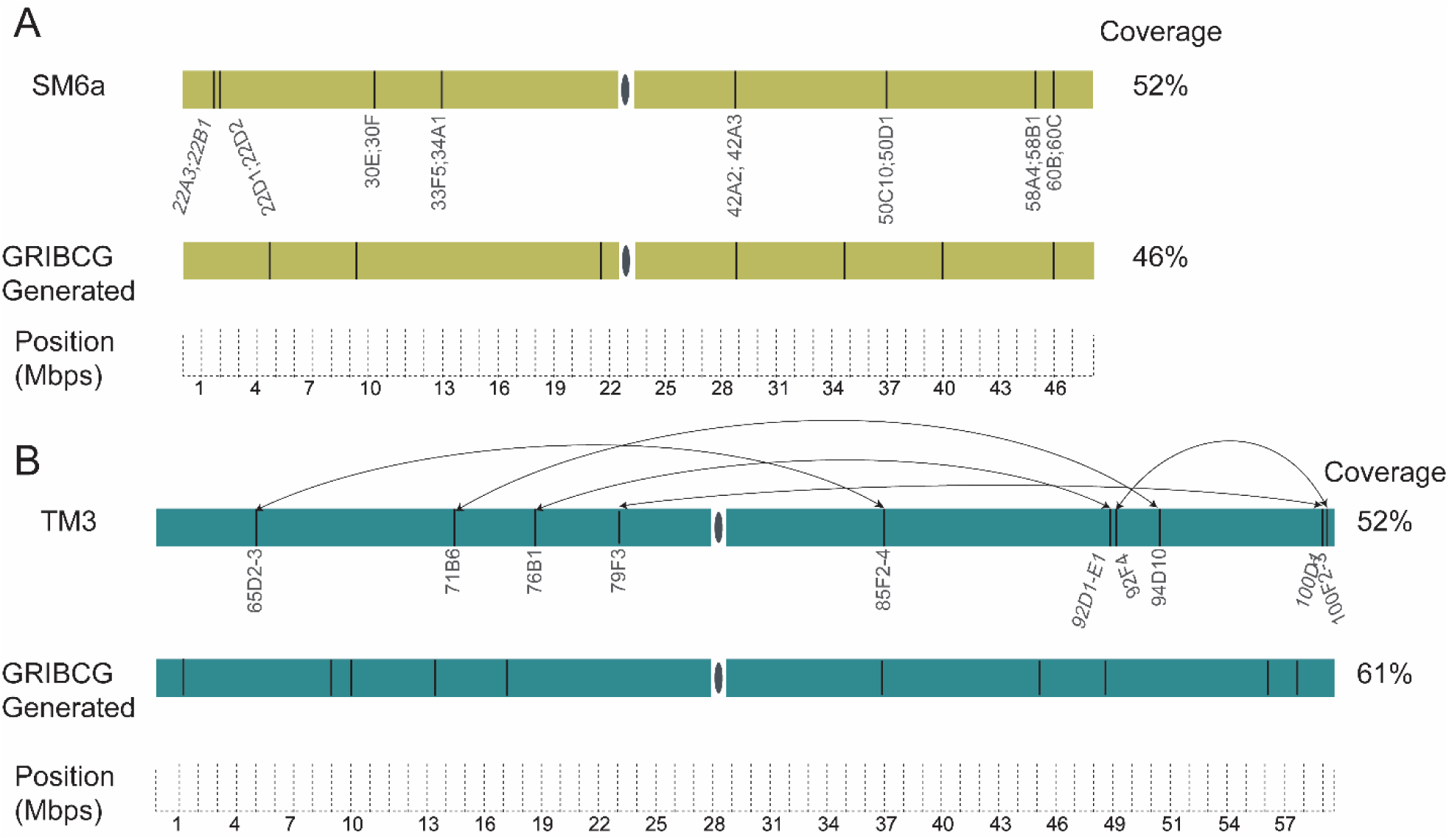
Comparison between GRIBCG results and existing fruit fly balancer chromosomes. A. *SM6a* is the most common second chromosome balancers in *D. melanogaster*, as described in Miller et al. 2016. Each vertical line represents a given double-stranded break. These breakpoints span the entire arm, encompassing both intergenic and genic regions. Below, the top ideal sgRNA for the same chromosome is depicted as generated by GRIBCG. B. *TM3* in the most common third chromosome balancers in *D. melanogaster.* Arrows indicate the breakpoints corresponding to the known sequence of inversions in the generation of *TM3*. Below, the top ideal sgRNA for the same chromosome is depicted as generated by GRIBCG.

## Conclusion

GRIBCG is a fast and easy-to-use tool for the selection of sgRNAs in the rational design of balancer chromosomes. While previous work has demonstrated successful generation of balanced regions in *C. elegans* and *Danio rerio* [5, 7], our tool is the first designed to create a completely balanced chromosome with the use of a single sgRNA. Experimentally, using a single sgRNA would eliminate the need for multiple rounds of transformation, and decrease the number of generations that need to be screened in order to identify a completely balanced chromosome. Thus, our work offers the possibility of expanding the use of balancer chromosomes to multicellular organisms other than *D. melanogaster.*

## Availability and requirements

Project name: GRIBCG

Homepage: https://sourceforge.net/p/gribcg/code/ci/master/tree/

Operating System: Linux (Ubuntu 18.04)

Programming Languages: Perl and R

License: none

No restrictions of use for academic or non-academic purposes.

## Abbreviations

Mbp: Megabase-pairs; GUI: Graphical User Interface; sgRNA: single-guide RNA; CFD: Cutting Frequency Determination; PTS: Potential Target Site; PSS: Partial Sequence Site; Gb: Gigabyte; Mb: Megabyte;

## Acknowledgements

We are grateful to Dr. Jung Choi, director of MS in Bioinformatics at Georgia Tech, for his valuable feedback on this project.

## Availability of data and materials

GRIBCG is available from SourceForge at: https://sourceforge.net/p/gribcg/code/ci/master/tree/

## Genome versions utilized for this work

*O. sativa* genome: ftp://ftp.ncbi.nlm.nih.gov/genomes/all/GCF/000/005/425/GCF_000005425.2_Build_4.0/GCF_000005425.2_Build_4.0_genomic.fna.gz

*A. thaliana* genome: ftp://ftp.ncbi.nlm.nih.gov/genomes/all/GCF/000/001/735/GCF_000001735.4_TAIR10.1/GCF_000001735.4_TAIR10.1_genomic.fna.gz

*C. elegans* genome:ftp://ftp.ncbi.nlm.nih.gov/genomes/all/GCF/000/002/985/GCF_000002985.6_WBcel235/GCF_000002985.6_WBcel235_genomic.fna.gz

*M. musculus* genome:ftp://ftp.ncbi.nlm.nih.gov/genomes/all/GCF/000/001/635/GCF_000001635.26_GRCm38.p6/GCF_000001635.26_GRCm38.p6_genomic.fna.gz

*D. rerio genome:*ftp://ftp.ncbi.nlm.nih.gov/genomes/all/GCF/000/002/035/GCF_000002035.6_GRCz11/GCF_000002035.6_GRCz11_genomic.fna.gz

## Authors’ contributions

B.B.M. developed the algorithms and coded the software package. B.B.M. and L.S.C. design the software and analyzed results. All authors read and approved the final manuscript.

## Ethics approval and consent to participate

Not applicable.

## Consent for publication

Not applicable

